# Deep learning of the splicing (epi)genetic code reveals a novel candidate mechanism linking histone modifications to ESC fate decision

**DOI:** 10.1101/189183

**Authors:** Yungang Xu, Yongcui Wang, Jiesi Luo, Weiling Zhao, Xiaobo Zhou

**Affiliations:** Center for Bioinformatics and Systems Biology, Wake Forest School of Medicine, Winston-Salem, NC, 27157, USA; Key Laboratory of Adaptation and Evolution of Plateau Biota, Northwest Institute of Plateau Biology, Chinese Academy of Sciences, Xining, Qinghai, 810008, China; Center for Systems Medicine, School of Medical Bioinformatics, University of Texas Health Science Center at Houston, TX, 77030, USA

**Author notes:** These authors should be regarded as joint first authors (Yungang Xu, Yongcui Wang).

## Abstract

Alternative splicing (AS) is a genetically and epigenetically regulated pre-mRNA processing to increase transcriptome and proteome diversity. Comprehensively decoding these regulatory mechanisms holds promise in getting deeper insights into a variety of biological contexts involving in AS, such as development and diseases. We assembled splicing (epi)genetic code, DeepCode, for human embryonic stem cell (hESC) differentiation by integrating heterogeneous features of genomic sequences, 16 histone modifications with a multi-label deep neural network. With the advantages of epigenetic features, DeepCode significantly improves the performance in predicting the splicing patterns and their changes during hESC differentiation. Meanwhile, DeepCode reveals the superiority of epigenomic features and their dominant roles in decoding AS patterns, highlighting the necessity of including the epigenetic properties when assembling a more comprehensive splicing code. Moreover, DeepCode allows the robust predictions across cell lineages and datasets. Especially, we identified a putative H3K36me3-regulated AS event leading to a nonsense-mediated mRNA decay of BARD1. Reduced BARD1 expression results in the attenuation of ATM/ATR signalling activities and further the hESC differentiation. These results suggest a novel candidate mechanism linking histone modifications to hESC fate decision. In addition, when trained in different contexts, DeepCode can be expanded to a variety of biological and biomedical fields.

## INTRODUCTION

Alternative splicing (AS) is one of the most important precursor (pre-) mRNA processing (1) to increase the mRNA- and protein-diversity in tissue- and development-dependent manners (2). Its mechanisms include alternative promoters, preferential usage of exons or splice sites, scrambling of exon order, and alternative polyadenylation (3). A number of genome-wide studies have revealed its prevalence in human (1,2), yeast (4), worms (5,6), and flies (7). AS is an integral part of differentiation and developmental programs and contributes to cell lineage and tissue identity. Aberrant splicing may result in developmental abnormalities, hereditary diseases (8) and cancers (9). Therefore, the faithful regulation of AS is essential for providing specific characteristics of cells and tissues, and for their responses to internal and external environmental changes (10).

AS regulation has long been thought to involve mostly RNA-binding proteins, including splicing factors (SF) and auxiliary proteins, that bind pre-mRNAs near variably used splice sites (SS) and modulate the efficiency of their recognition by the basal splicing machinery (the spliceosome). Large-scale quantification of AS combined with genome-wide identification of *in vivo* binding sites of splicing regulators provide an unprecedented global view of splicing regulatory networks (11) and enable the predictions of AS outcomes based on these genetic attributes (12). Given the conceptions from semiotics, the increasing variety and interactive properties of both genetic and epigenetic features has led to the use of the term ‘code’ to describe the predictive and heritable (epi)genetic attributes that specify patterns of gene expression through differentiation and development. Likewise, in the context of RNA splicing, combining the *trans-acting* regulators with *cis-acting* RNA elements have enabled the inference of splicing genetic code (13), which revealed a number of novel regulators critical for embryonic stem cell (ESC) differentiation, such as MBNL (14) and SON (15). However, these genetic controls have been found far from sufficient to explain the whole picture of AS (16) and the static nature of genome sequence limits the ability to generate cell-type-specific predictions for samples not previously used for training (17).

It is clear that a splicing code must incorporate various features that act together to decode the splicing at multi-levels from chromatin to RNAs *per se*. Furthermore, a code should be able to reliably predict the regulatory properties of previously uncharacterized exons and the effects of both genetic and epigenetic variations within the regulatory elements. As expected, genome-wide association studies have revealed the global correlations between histone modifications (HMs) and AS events (18-22). More specifically, AS is also under controls of the epigenetic mechanisms due to its co-transcriptional occurrence (23). Especially, the HMs, which define the chromatin states, determine not only what parts of the genome are expressed but also how they are spliced. For example, H3K4me3 (24), H3K9me3 (25), H3K36me3 (26,27), and acetylation of H3 (28) are emerging as major regulators of AS, either by directly recruiting splicing factors (SFs) or indirectly modulating transcriptional elongation rate (29). These emerging findings have opened a new avenue and are promising to decipher a more extended splicing code, by which we may address these imperfections. Therefore, our goals are to decipher the extended splicing (epi)genetic code compiling not only the genetic elements but also the epigenetic properties, to understand how they interplay with each another in regulating AS and contributing to various biological processes, such as development and disease progress.

The increasingly rapid growth in the volume of ‘multi-omics’ data (e.g. genomics, epigenomics, transcriptomics) makes it possible to achieve these goals, yet requires more powerful computational approaches to handle and draw biological mechanisms from these multi-omics datasets. Machine learning algorithms, such as *k*-means clustering, support vector machines (SVM), and random forests (RF) have been applied to these omics data, separately (30). Recently, deep learning has gained attentions due to its high performance and generalized characteristics for analyzing complex data of various contexts (31,32), such as image and speech recognition (33-35), natural language understanding (arXiv preprint arXiv:1409.0473 and arXiv:1603.01417), and most recently computational biology (12,36-39). It aims to replace handcrafted features with efficient algorithms for unsupervised or semi-supervised feature learning and hierarchical feature representation using a cascade of nonlinear processing unit. Our previous work (32) reveals its potential to capture the latent structures of cancer patients and enable the generation of distinct survival subtypes of cancer patients with unique clinical and molecular characteristics based on those latent structures. Thus, deep learning can also be used to efficiently decipher the splicing (epi)genetic codes involving heterogeneous genetic and epigenetic factors. To this end, we implemented a deep neural network, DeepCode (Figure 1), which took both genomic sequences and epigenetic features as inputs to predict the splicing patterns in a variety of contexts, including but not limited to development and disease.

**Figure 1.**
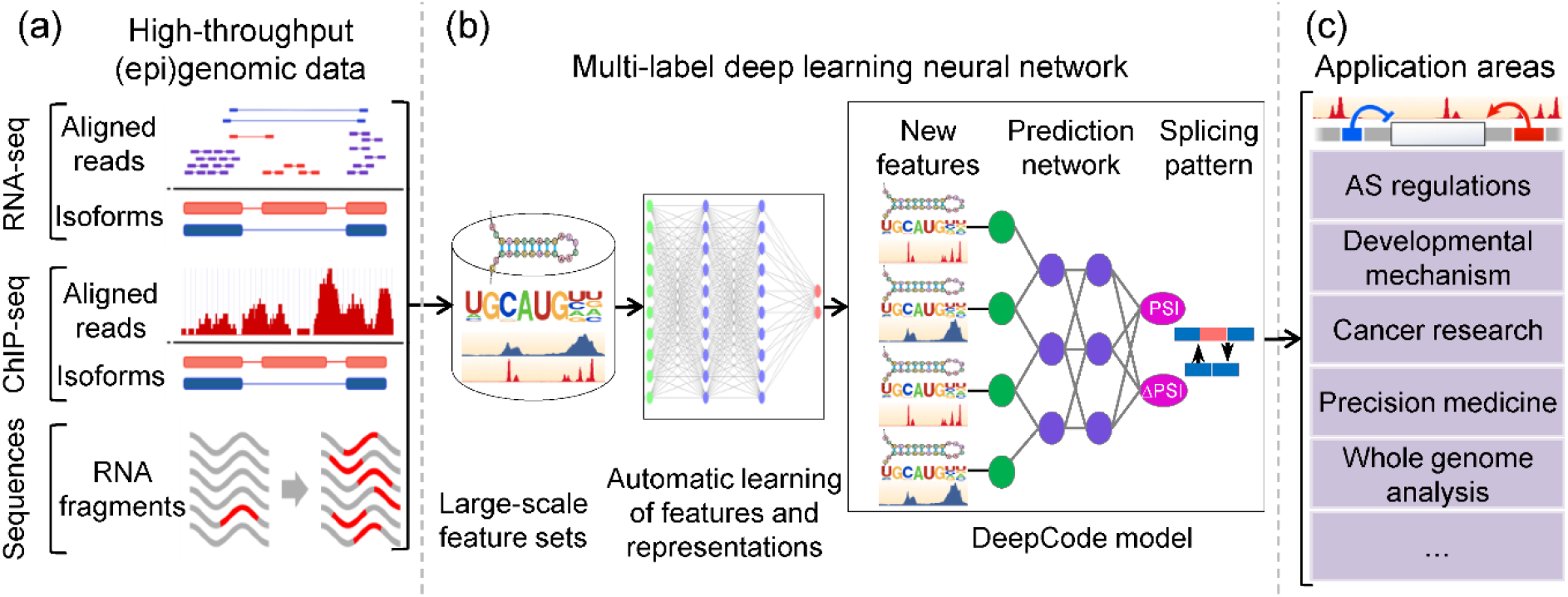
Schematic view of DeepCode. **(a)** Multiple high-throughput data are integrated to feed the deep learning neural network. RNA-seq data are used for AS events identification and ‘percent spliced in’ (PSI) quantification; ChIP-seq data for epigenomic features; and RNA sequences for genomic features. **(b)** A multi-label deep neural network is proposed to learn the (epi)genetic code for AS. The code will be predictive for inclusion levels (PSIs) and their changes (ΔPSI) upon hESC differentiations. **(c)** The resulting model, DeepCode, can then be used to investigate AS regulations genetically and epigenetically, and further to evaluate effects of genetic and epigenetic variants on AS during normal development, disease progressions, and so on.

ESCs are pluripotent cells that can proliferate indefinitely while retaining the capacity to differentiate into multiple lineages of three germ layers and provide a vital tool for studying the regulation of early embryonic development and cell fate decision (40). A number of genome-wide studies have made remarkable progress in understanding the ESC fate decision. On one hand, AS provides a powerful mechanism to control developmental decisions in ESCs (15,41,42) and undergoes extensive controls from both genetic and epigenetic mechanisms (23). On the other hand, HMs plays crucial roles in ESC maintenance and differentiation (43) by determining not only what parts of the genome are expressed, but also how they are spliced. It is intuitive to raise and dedicate to answer the questions of whether and how the HMs contribute to splicing code and collectively affect cell differentiation.

Therefore, in this work, we integrate multi-omics data of hESC differentiation with the proposed deep learning approach, DeepCode, to decipher an extended splicing code for ESC fate decision. With the advantages of epigenetic features, DeepCode significantly improves the performance in predicting splicing patterns during hESC differentiation. In addition, epigenetic properties outperform genetic elements and play dominant roles in decoding AS outcomes. We also found that DeepCode is useful for understanding the (epi)genetic determinants of embryonic development by revealing a novel candidate mechanism linking HMs to ESC fate decision. In addition, DeepCode is designed for its scalability in data types and extensibility in applications, thus holds the potential applications in a variety of biological and biomedical fields when trained in different contexts.

## MATERIAL AND METHODS

### Datasets

To decipher the splicing (epi)genetic code for embryonic development, we used two datasets from independent research groups for primary model training and evaluations. The first dataset was from four cell lineages directly derived from hESCs (H1 cells), including trophoblast-like cells (TBL), mesendoderm (ME), neural progenitor cells (NPC), and mesenchymal stem cells (MSC), which was generated from Bin Ren’s lab (43). We also included IMR90 for Ren’s dataset, a cell line for primary human fetal lung fibroblasts, as an example of terminally differentiated cells. The second dataset was from Alexander Meissner’s lab (44) with regard to a population of cells derived directly from hESCs (HUES64) representing each of three embryonic germ layers, including ectoderm (dEC), mesoderm (dEM), and endoderm (dEN). Matched ChIP-seq data of 16 HMs and DNase, and RNA-seq data (two replicates) for each cell type were collected and analyzed (Supplementary Methods, M1). In this study, we mainly focused on two types of mostly occurred AS events, mutually exclusive exons (MXE) and skipped exons (SE). These AS events were identified from the RNA-seq data with the ‘per spliced in’ changes |ΔPSI| > 0.1 at FDR < 0.05, based on measurements used by rMATS (45) (Supplementary methods M1). We defined these AS events as hESC differentiation-related AS events, including 3,513 MXE and 3,678 SE events for Ren’s dataset (Supplementary Figure S1a, Table S1, and Dataset D1), which were the primary dataset for model training and evaluation. We also identified the AS events for Meissner’s dataset (Supplementary Table S1), which were used as the independent dataset for evaluations of model robustness. The genomic and epigenomic features were learned from the ±150bp regions surrounding both the 5’ and 3’ splice sites (SS) of the AS exons (Supplementary Figure S1b). We identified 136 and 68 epigenomic features of 17 ChIP-seq datasets for SE and MXE exons, respectively (Supplementary Methods M1 and Table S1). Besides, 421 pre-defined sequence features are also considered as genomic features, including known motifs, new motifs, short motifs and transcript structures, which were assembled as splicing genetic code and originally described in reference (13). Additionally, we also employed the extensive resources from Roadmap (46,47) and ENCODE (48,49) projects, wherein 5 ESC or iPSC, 8 ESC-derived cell types, and 43 adult cells/tissues were considered (Supplementary Dataset D2 and Methods M5).

### The DeepCode

DeepCode, an extended splicing (epi)genetic code, is proposed to predict how given exons are spliced differentially in different cell lineages derived from hESCs. The method takes multi-omics data (RNA-seq, ChIP-seq, and RNA sequences) as inputs for large-scale feature learnings and representations (Figure 1a). A multi-label deep neural network is then automatically trained to learn and integrate multi-omics features; the trained model (DeepCode) enables the prediction of splicing patterns represented as inclusion levels (PSIs) and their changes (ΔPSIs) in different conditions (Figure 1b). Trained in different contexts, DeepCode can be used in not only understanding AS regulations, but also a variety of biological and biomedical topics, including but not limited to developmental mechanisms and cancer researches (Figure 1c).

#### Defining the learning tasks

Because of the position-dependent biases in the coverage of RNA-seq reads (50), estimated exon inclusion levels (i.e., PSIs) may not be accurate enough for model training across samples (cell lineages). To avoid directly learning the absolute values of PSIs, we discretized the inclusion levels into High (H), Medium (M), and Low (L), based on the global distribution of PSIs (Supplementary Methods M2 and Figure S2). In addition, in order to extend the model prediction capability for lineage-specificity, we also applied the model to predict PSI changes (ΔPSIs) upon hESC differentiation. In short, we had two learning tasks. The first one is to predict the splicing outcome of a given exon in a specific cell lineage; and the second to predict the ΔPSIs induced by differentiation, represented as either increased (I) or decreased (D). We assembled these tasks into five-dimensional binary vectors, <H, M, L, I, D>, and named it as splicing patterns (SPs, Supplementary Figure S3 and Methods M2). This definition makes the learning tasks a multi-label learning problem (MLP). Usually, there are two strategies for MLP - either transforming MLP as several binary classification problems, or transforming MLP into a multi-class classification problem (51). We used the former one adapted from previous work (52) to learn the SPs. Specifically, to learn the SPs represented as a five-dimensional binary vectors, five binary classifiers were introduced to distinguish exons with High inclusion level from others, with Medium inclusion level from others, with Low inclusion levels from others, with Increased inclusion level from the others induced by differentiation, and with Decreased inclusion level from others induced by differentiation, respectively.

#### Designing the architecture and mathematical representation of the DeepCode

Previous work only introduced sequence features to decode SPs (13,52). Given the emerging crucial roles of epigenetics in AS regulation, we also introduced epigenomic features for a more comprehensive decoding of SPs. To integrate both sequence and epigenomic features, DeepCode was designed as a multi-layer neural network, which includes transformation layers for feature abstraction and a fusion layer for feature integration (Supplementary Figure S3). Furthermore, the multi-label architecture was introduced to decode the SPs represented as five-dimension binary vectors, which allows learning shared features between outputs, thereby improving generalization performance, and markedly reducing the computational cost of model training compared to learning independent models for each task (13).

In the term of SP prediction based on integration of genomic and epigenomic data, the meaningful features can be individual sequence motifs or *k*-mers and epigenetic signals at the lowest layers, combined motifs and epigenetic signals at the intermediate layers, and complex (epi)genomic code at the highest layers. The specially designed DeepCode allows integrating diverse input feature types and is able to capture both inner- and inter-feature relationships through its transformation and fusion layers. Suppose we have a total of exons with a total of epi(genomic) feature types and layers in a multilayer neural network (Supplementary Figure S3). Denote 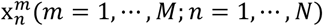 *as* the original value of inputted feature associated with *m*th epi(genomic) feature type in the th exon; 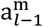 and 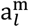 as the input and output of the *l*th layer for *m*th feature type, *l* = 1,*E* (*E*, the number of transformation layers); and *a_t-1_* and *a_t_* as the input and output of *t*th fusion layer, **t* = *E*+1,…,*L*−1*. Let 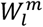 and 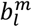 as the weight matrix and bias vector of the transition function from (*l* - 1)th to *l*th transformation layer, respectively, and *W_t_* and *b_t_* as the weight matrix and bias vector of the transition function from (*t* - 1)th to *t*th fusion layer, respectively. Given exon associated with the *m*th feature type x*^m^*, the transition function from the (*l*-1)th to the *l*th transformation layer, and (*t*-1)th to *t*th fusion layer were calculated as follows:

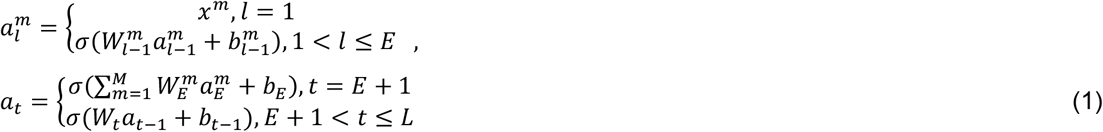

where σ(·) is sigmoid function, defined as 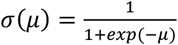.

Let 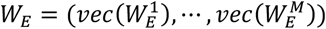, where 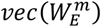 is obtained from matrix 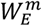 by stacking the columns of 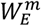 one on top of another, *m* = 1,…, *M*. Then the optimal weights for each layer can be obtained by the following optimization problem:

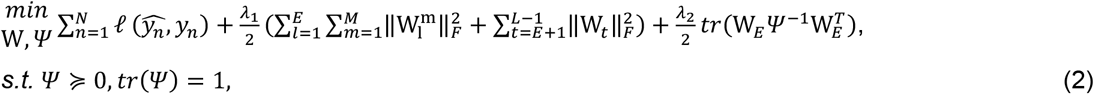

where *l* is the loss function measuring the discrepancy between the observed SP y = (y_1_,…*,y_N_*) and the estimated SP 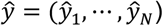. The regularized least square (RLS) was introduced as the loss function to access the discrepancy between the observed SPs and the estimated ones. *λ*_1_ and *λ_2_* are regularization parameters; positive semi-definite matrix *ψ* ∈ **R*^×*M*^* models the inter-feature relationship; *tr*(*ψ*) *=* 1 is applied to restrict complexity of the model, as suggested in (52), and ||*w*||*_F_* is Frobenius norm of matrix *W*. The first two regularization terms in the optimal function are used for sparsity control and last one for managing the diversity among different feature types.

### Model implementation and parameters selection

Our previously proposed optimization method (32) were applied to iteratively minimize the optimization problem (Formula 2) with respect of weight matrix *W* and diversity control matrix *ψ*. By fixing *ψ*, the optimization problem (Formula 2) becomes an unconstrained optimization problem with respect of both vector and sparse matrix. The Frobenius norm of a matrix *W* is the 2-norm of its vector transition *vec*(*W*), which is obtained from matrix *W* by stacking the columns of *W* one on top of another. The positive semi-definite matrix *ψ* could be decomposed as a product of matrix *U* and its transpose 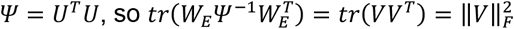, where *V = W_E_U^-1^* Then the Formula 2 with a fixed *ψ* becames a nonlinear unconstrained optimization problem with only respect to the vector, which was solved by “fminunc” function in MATLAB. When the optimized weight matrices 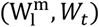 were obtained, the optimized *ψ* was obtained by introducing Cauchy-Schwarz inequality into a simple semi-definite optimization problem. The model parameters *λ*_1_*, λ_2_* were selected by five-fold cross-validation procedure, and the number of total layers in the network was predefined as four to facilitate rapid convergence and reduce over-fitting. The sparsity control for parameters was added in our optimization. It would force many of them close to zero and generate few of parameters with large values that indicated the (epi)genomic features strongly associated with splicing outcomes.

### Feature selection and (epi)genetic code assembling

The optimal weight matrices 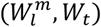 indicate the most contributed (epi)genomic features during SPs learning. Supplementary Figure S4 depicts the flowchart for feature selection after optimization problem (Formula 2). Suppose the multi-layer neural network only contains four layers, including input, transformation, fusion and output layers, the most contributed fusion notes were found based on the value of *W_L_*. The weight matrices for top fusion notes (*W_t_*) provide the evidence for determining the most contributed (epi)genomic feature types during the learning. The (epi)genomic variables that could be used as important features were traced based on the weight matrix of transformation layer 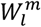. The candidate features were selected based on their importance that were calculated by multiplying their associating weights and used to evaluate their contributions to the splicing (epi)genetic code. The selected important features were evaluated by how the performance changed based on the selected features. The top important (epi)genomic features (150 in this study) were then assembled as the final splicing (epi)genetic code.

### Model evaluation and statistical analyses

Since the imbalance problem was raised due to one-versus-the-rest procedure, which was employed to implement MLP, and the precision-recall curve is a better index for evaluation the imbalance problem (13), the area under the precision-recall curve (AUPR) was introduced as the criteria to display the performance of the learning model. Evaluations were performed among different feature types across different cell lineage and datasets. We also compared the performances of DeepCode with other counterpart models. Specifically, two types of cross-validation strategies and one independent dataset were explored to test the model’s robustness and ability of predicting between cell lineages and datasets (Supplementary methods M3). The statistical significances were tested using Student’s t-test (R package) for Figures 2d, e, 3, 4c and S6c. Especially for Figures 2d and 2e, the tests were performed based on the y-axis values along with same x-axis values. Pearson correlation tests were used for 157 Figures 5, S8, S9, and S14.

**Figure 2.**
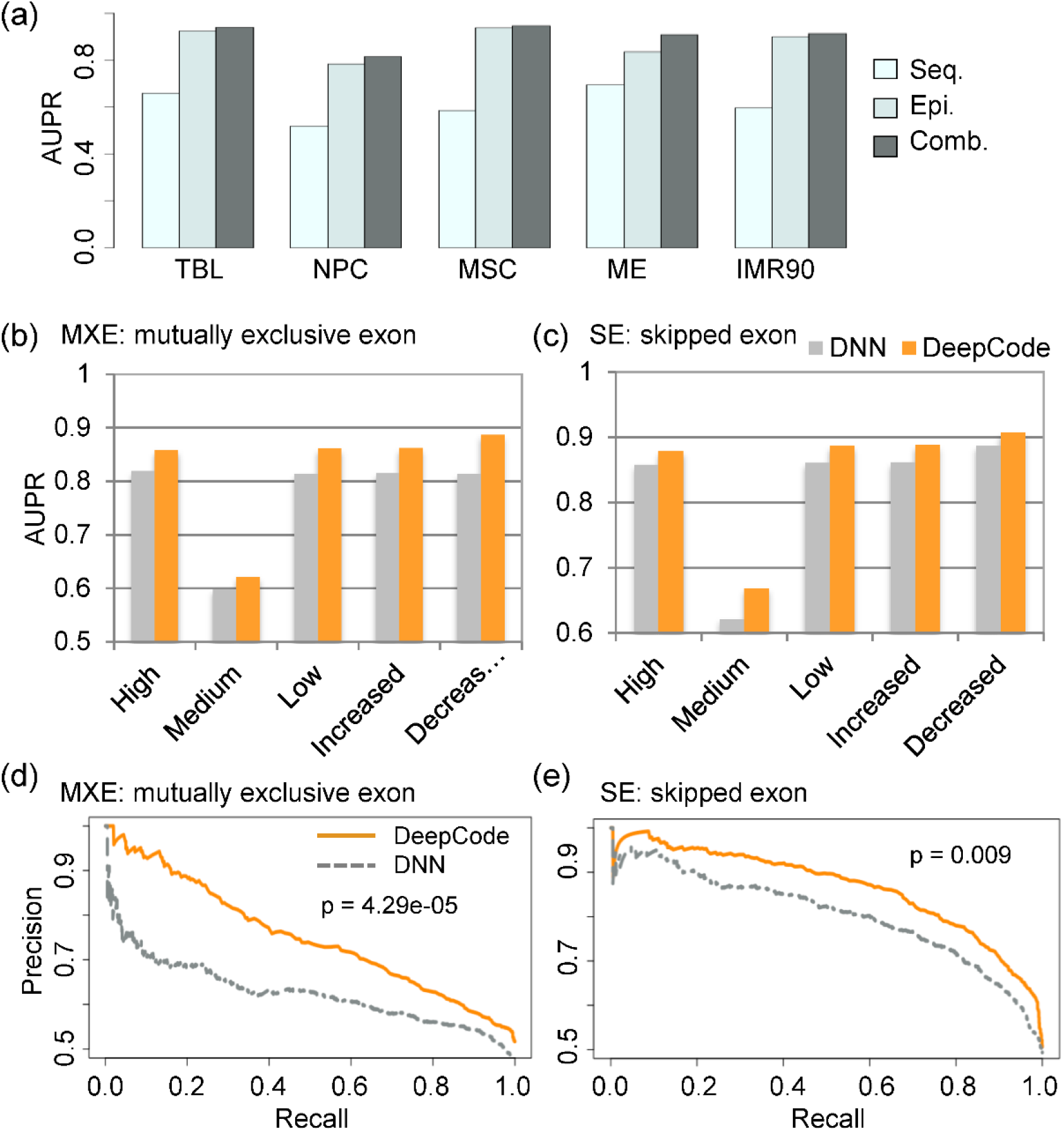
DeepCode improves the performance in decoding splicing patterns. **(a)** Epigenomic features improve model performance sharply, suggesting they are more important in decoding splicing patterns. Seq, sequence features alone; Epi, epigenomic features alone; Comb, combining the Seq and Epi together. **(b** and **c)** DeepCode improves performance for decoding splicing patterns for both MXE (b) and SE (c) events. Shown as the average AUPR across five cell lineages. **(d** and **e)** The precision-recall curves of predicting inclusion level changes for MXE (d) and SE (e) events during hESC differentiations. p, p-values based on Student’s t-test.

**Figure 3.**
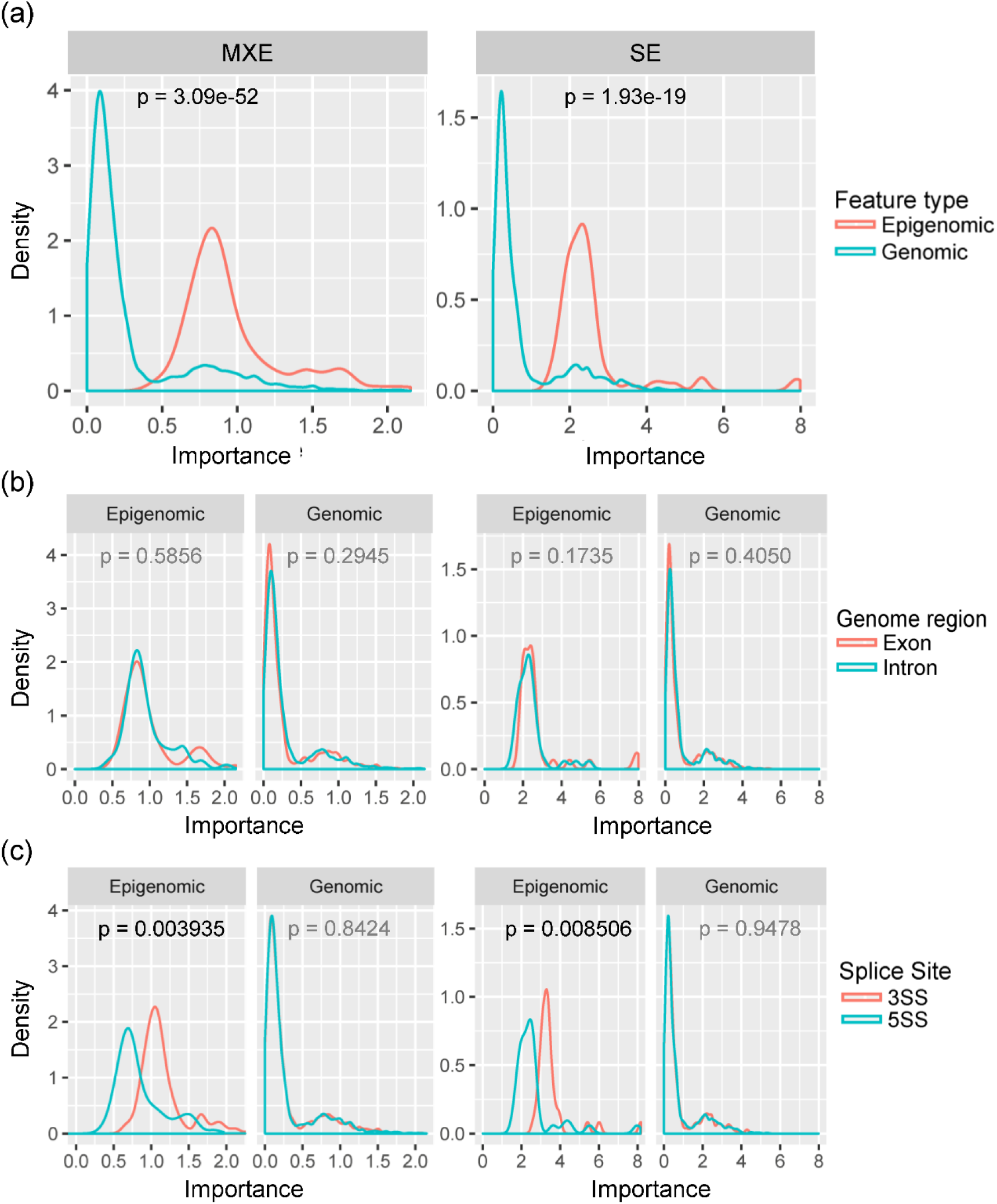
DeepCode reveals different aspects of the contributions of (epi)genomic features. **(a)** Epigenomic features are more important in deciphering splicing code. **(b)** Both genomic and epigenomic features from exonic and intronic regions show no difference in contributing to splicing code. **(c)** Epigenomic features around 3’SS have higher importance than around 5’SS; while genomic features are not significantly different. p, p-values based on Student’s t-test.

**Figure 4.**
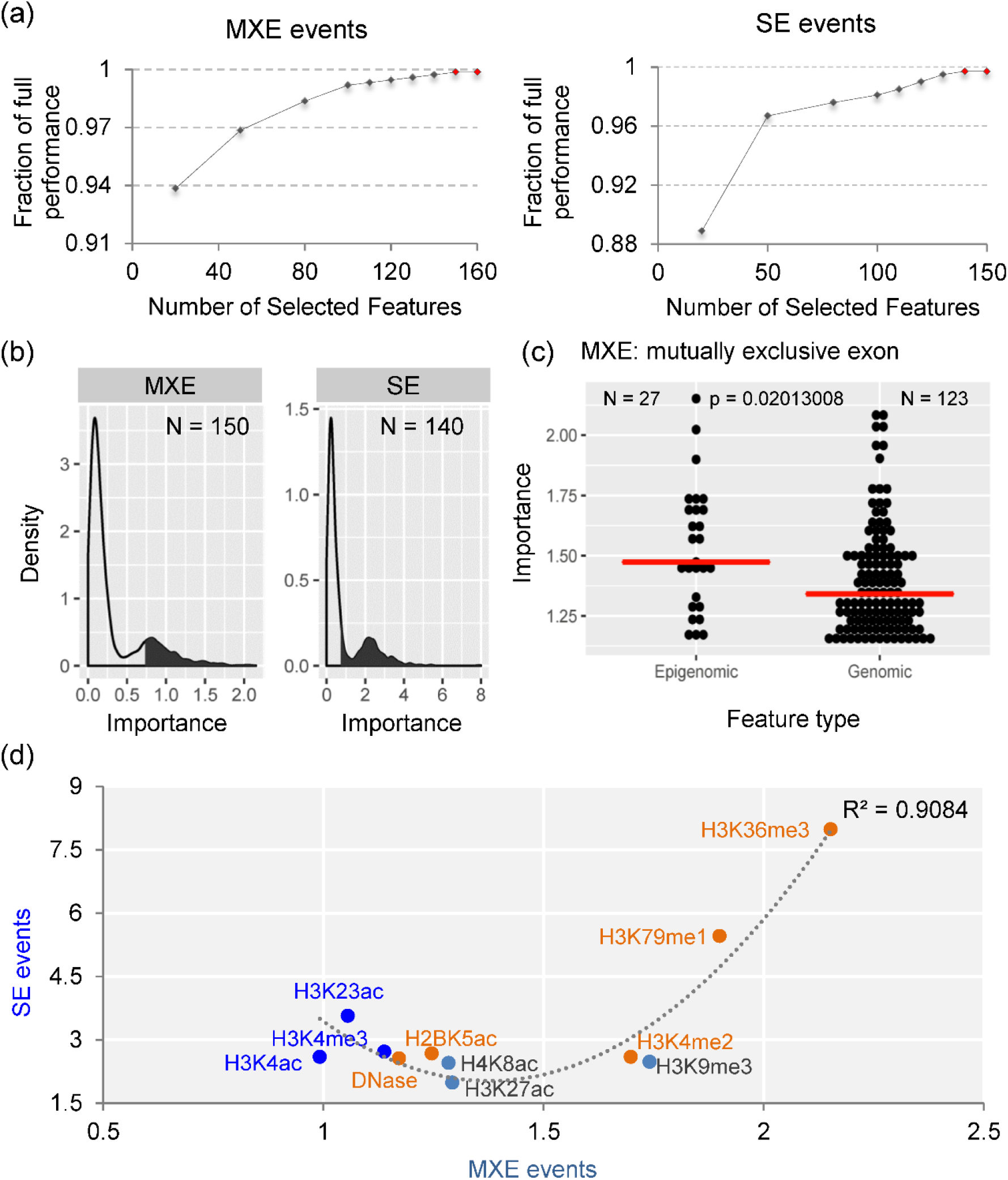
Top important features and DeepCode assembly. **(a)** The predictive performances of DeepCode increase with the number of top important features for MXE (left) and SE (right) events. **(b)** The distributions of feature importance and top important features (filled parts) for MXE (left) and SE (right) events. **(c)** The top important epigenomic features are consistently of higher importance than top important genomic features. Only the case of MXE events is shown and the case of SE events is provided in Supplementary Figure S6c. N, the number of involved top features; p, p-values based on Student’s t-test. **(d)** The top important epigenomic features are concordant between MXE and SE event, and consistent with the previous reports. Orange dots are the features shared by the splicing codes for MXE and SE events. R^2^ indicates the fitness of polynomial curve.

## RESULTS

We trained the primary DeepCode model based on the two types of AS events (MXE and SE) that occur during hESC differentiation to multiple cell lineages (Ren’s dataset). Various evaluations were employed to assess the performance of DeepCode. We show that DeepCode surpasses other state-of-the-art approaches in predicting how the transcripts will be spliced during multiple-lineage differentiations of hESCs and highlights the contributions of epigenomic features to decoding the SPs. DeepCode also facilitates getting deeper insights into the involvement of HMs in ESC fate decision through the regulation of AS.

### DeepCode improves the performance of decoding the splicing patterns

We first assessed the effects of (epi)genomic features on predicting performance. As shown in Figure 2a, the epigenomic features are more predictive than the genomic sequence features, and the combination of both improves performance much more than sequence features alone. This result suggests that epigenomic features play important roles in deciphering the splicing code of higher predictive power. We then compared our results with a commonly used deep learning framework, the deep neural network (DNN), which has been proven to outperform the Bayesian neural network and multinomial logistic regression model in learning the splicing genetic code (12). We implemented the DNN on Ren’s MXE and SE AS events across all cell lineages, and compared the AUPRs. DeepCode achieves higher AUPRs on both MXE and SE events, indicating that our approach outperforms DNN in both types of AS events (Figures 2b and 2c, Supplementary Tables S2 and S3).

We further validated the ability of DeepCode and DNN in predicting the inclusion level changes (increased or decreased), which directly reflects the lineage-specificity. Both methods were performed on the Ren’s MXE and SE events across all five cell lineages to test their ability of distinguishing the lineage-specificity. DeepCode has superior performance (Figures 2d and 2e) over the DNN. It is because the multi-layer neural network of DeepCode is specially designed for multi-omics data integration, and also due to contribution of epigenomic features. Taken together, our analyses showed that DeepCode improves the performance of decoding the splicing patterns and highlights the importance of epigenomic features.

### Epigenetic properties outperform genetic elements in decoding the splicing patterns

The contributions of features in decoding SPs can be uncovered through the weight matrices in the multi-layer neural network and were measured as ‘importance’ (see Material and methods). The features of higher importance contribute more to splicing code. Taking into account all features (3504 features for MEX and 1752 features for SE), we observed that epigenomic features are more important than genomic ones (Figure 3a and Supplementary Figure S5a). It strongly suggests that the epigenetic properties, such as histone modifications, should be taken into account when assembling a more accurate splicing code. We then compared the features with respect of their genomic locations. Both genomic and epigenomic features show no difference in importance between exonic and intronic regions (Figure 3b and Supplementary Figure S5b), indicating that AS regulatory machinery does not prefer to exonic or intronic regions around the splice sites (SS). We further compared features regarding to the 3’SS or 5’SS. In this regard, epigenomic features around 3’SS are more important than around 5’SS, while genomic features show no significant difference between them (Figure 3c and Supplementary Figure S5c). In addition, epigenomic features display more diversity in importance among different feature regions (Supplementary Figure S6a). For MXE events, though both genomic and epigenomic features showed no significant preference of importance between the upstream and downstream exons, a moderate difference was observed for epigenomic features (Supplementary Figure S6b). Taken together, we showed that epigenomic features are distinct in many aspects with the genomic features and, especially, epigenomic features outperform genomic ones in decoding the SPs.

### Assembling a splicing (epi)genetic code

We aimed to assemble a code that is able to predict the splicing patterns of all AS exons as accurately as possible, based on RNA sequence features and HM ChIP-seq signals. To assemble such a code that encompasses the top important features to decode the SPs, we performed single layer neural network analyses on the recursively selected features and evaluated their importance through the caused changes in AUPRs with performing multi-layer neural network analyses on full features. For the MXE events, when the numbers of selected features reaches 150, adding more features does not further improve the performance (Figure 4a left panel). For the SE events, the minimum required features for achieving the highest performance is 140 (Figure 4a right panel). These results suggest that only a small portion (4% for MXE and 8% for SE) of features are sufficient to predict the SPs, accounting for up to 99.8% of the performances with all features included (Figure 4b). Therefore, we assembled the final (epi)genetic codes containing 150 features for both the MXE and SE events for consistency reason (Supplementary Figure S7). Consistent with the global profiles of full features, the top important epigenomic components display higher importance than the top genomic ones (Figure 4c and Supplementary Figure S6c). Moreover, the epigenetic components of the final splicing codes are concordant between MXE and SE events (Figure 4d) and consistent with the previous reports (21). For example, the top two epigenetic features (H3K36me3 and H3K79me1) have been reported to be mostly related to AS (21).

Genetic elements have been previously assembled as splicing genetic code (13). We further compared the DeepCode with this genetic code regarding to the correlations between the DeepCode importance and the genetic code weights (shown as ‘Barash *et al*. weight) of the same group of genomic features. We found that the genomic features are highly correlated in their importance (weight) between these two independent models in decoding the splicing patterns. It is consistent for the model trained on full features (Supplementary Figure S8) and finally assembled splicing code (Figure 5). We also observed some outlier features not correlated between these two models, which are reasonable upon the fact that epigenomic features play crucial roles and could abolish some redundant contributions of genomic features that got high weights in previously assembled code. All together, we assembled a splicing (epi)genetic code composed of 150 (epi)genomic features, of which the genetic components show high correlation with those of a previously assembled code (13).

**Figure 5.**
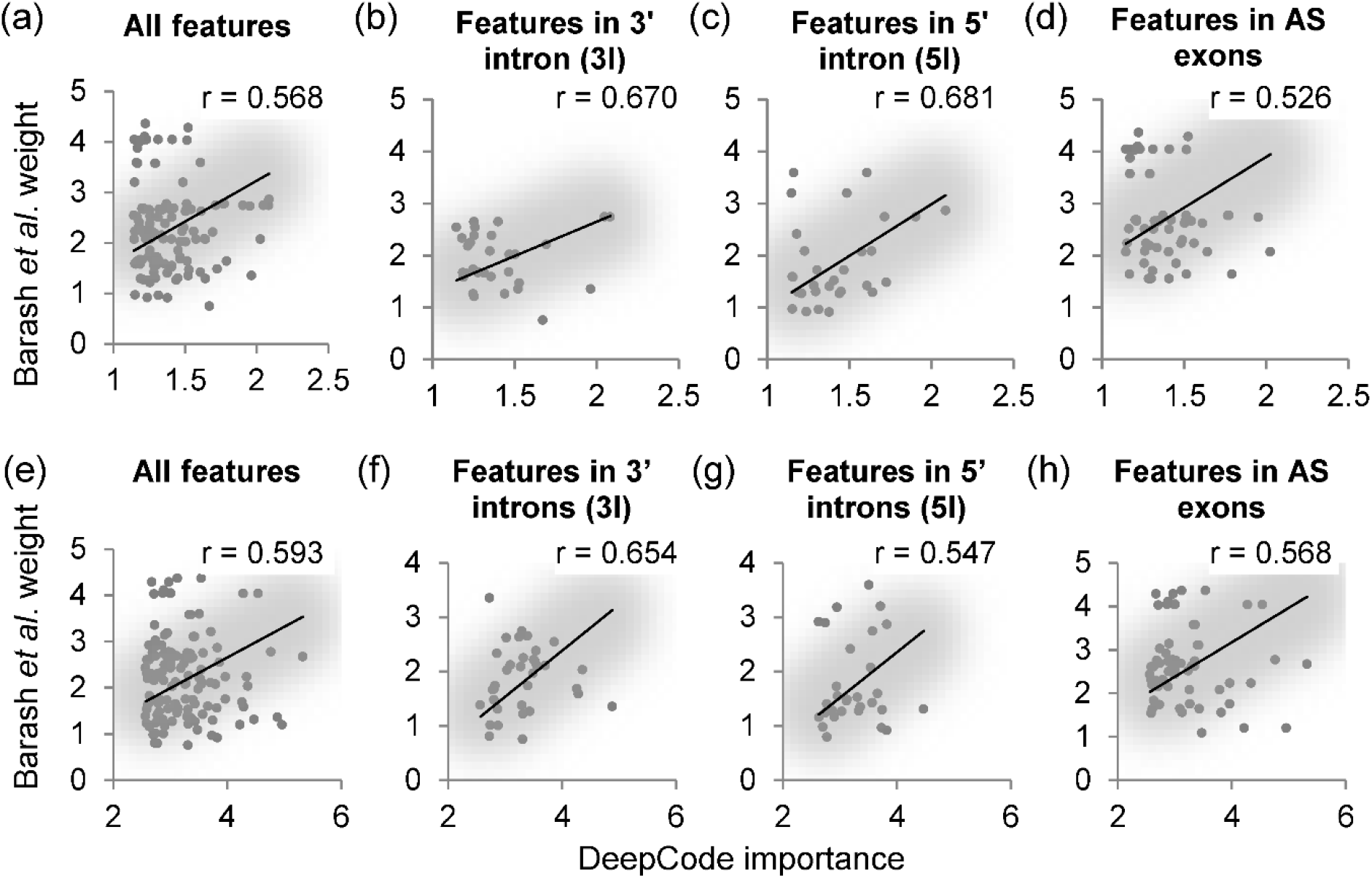
The assembled DeepCode shows high correlations with the previously assembled splicing genetic code regarding to genetic features’ importance. **(a-d)** The correlations for MXE events. Both upstream and downstream exons were pooled together. **(e-h)** The correlations for SE events. Black lines represent the linear fittings; r, the correlation based on Pearson correlation test.

### DeepCode allows the prediction of AS patterns across cell lineages and datasets

To test the robustness of the assembled splicing (epi)genetic code for prediction of AS patterns across developmental lineages and datasets, we employed independent dataset and implemented two additional evaluation strategies (Supplementary Methods M3).

Firstly, we trained DeepCode using AS events of two, three, and four of the five cell lineages from Ren’s dataset to predict those of the rest lineages, respectively (i.e. cross-lineage tests, shown as 2vs3, 3vs2, and 4vs1 in Supplementary Tables S4 and S5; each individual test is named as T1-T25). The performances were comparable with five-fold cross-validation (Supplementary Tables S4 and S5). We also checked whether the imbalanced sample sizes between training and testing datasets will affect the predicting performance. We only observed better predictions for some of those tests using cell lineages of larger sample sizes to predict the remaining cell lineages with smaller sample sizes, such as the tests T3-5, T12-15, and T22-25. It suggests that the sample imbalance across cell lineages has slight impact on performances of cross-lineage tests, especially for MXE events (Figures 6a, b, Supplementary Tables S4 and S5). Furthermore, the feature importance of cross-lineage tests is highly correlated with the importance based on five-fold cross-validations (Supplementary Figures S9 and S10). It suggests that the model trained based on some cell lineages is able to produce reliable predications for other cell lineages, which is supported by the observation that the changes of HMs during cell differentiation are lineage-independent, i.e., no lineage-specificity (Supplementary methods M4 and Figure S11).

**Figure 6.**
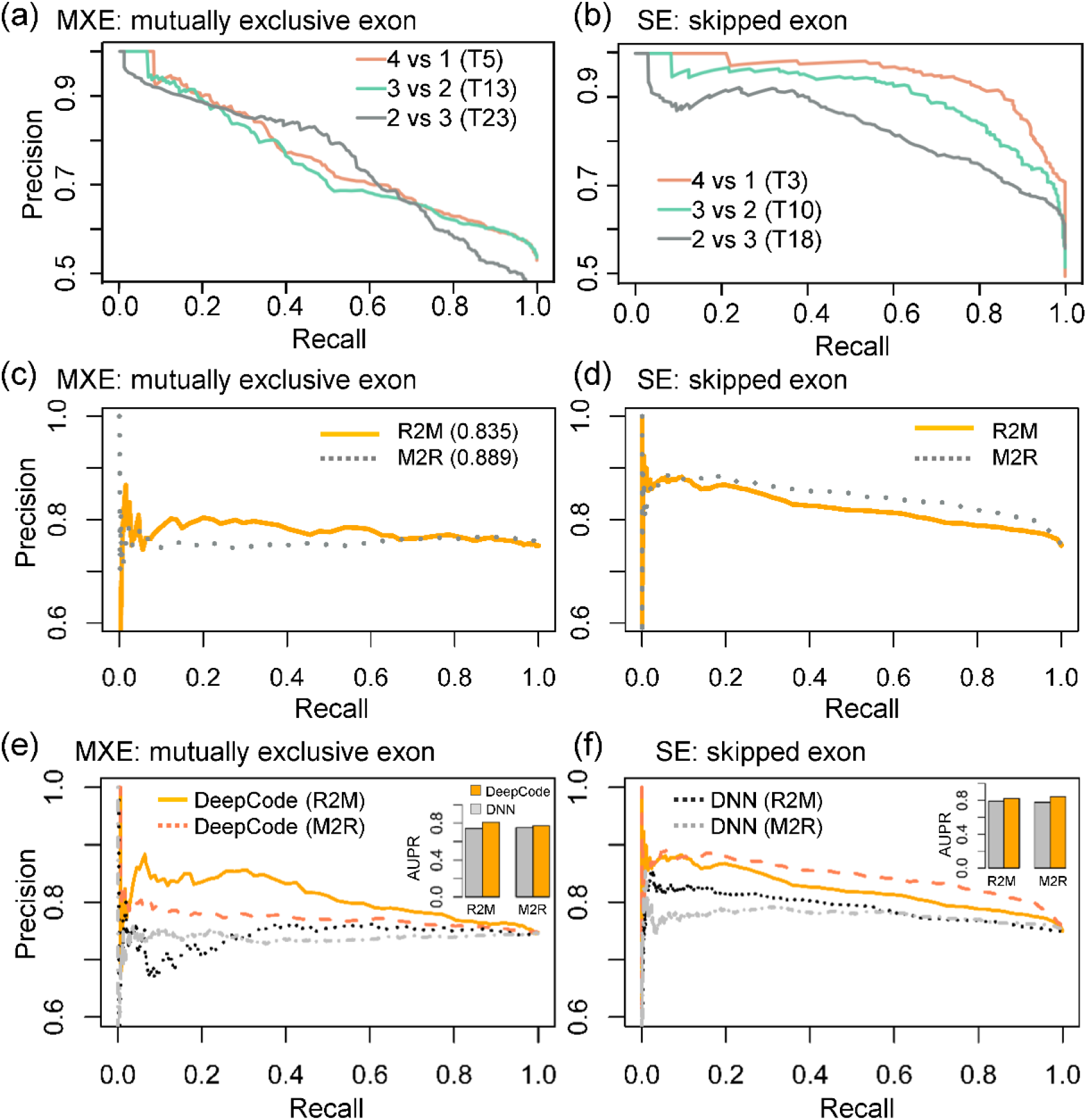
DeepCode allows the prediction of AS patterns across developmental lineages and datasets. (**a** and **b**) The precision-recall curves for the leave-p-out cross-validations (cross-lineage tests) of the highest AUPRs, based on the samples of MXE (a) and SE (b) events, respectively. (**c** and **d**) The precision-recall curves of cross-dataset tests for MXE (c) and SE (d) events, only the tests of the best AUPRs (Table S6-S9) are shown. (**e** and **f**) The performance comparisons between DeepCode and DNN for the cross-dataset tests. The insets show the AUPR of each test.

Secondly, we implemented independent tests between the Ren’s and Meissner’s (44) datasets to evaluate the robustness of DeepCode for cross-dataset predictions (Supplementary Methods M3). We tested the performances of DeepCode in predicting AS patterns between these two datasets reciprocally, i.e. DeepCode was trained using one dataset to predict the other one. The AUPRs ranged from 0.71 to 0.89, some of which are even better than the cross-lineage tests (Figures 6c, 6d, and Supplementary Tables S6-S9). No matter which dataset was used for training and the other one for testing, the DeepCode reaches nearly the same performances. Meanwhile, DeepCode holds better performances than DNN in cross-dataset tests (Figures 6e and 6f). Altogether, DeepCode allows the prediction of AS patterns among cell lineages and is robust across datasets.

### DeepCode reveals a novel candidate mechanism linking histone modifications to fate decision

Following the insights gained from DeepCode, we found that a number of core component genes of the ‘ATM/ATR-mediated DNA damage response’ pathway undergo H3K36me3-associaated AS during hESC differentiation (Supplementary Figures S12a and S13). This pathway is activated in S/G2 phases and has recently been reported as a gatekeeper of the pluripotency state dissolution (PSD), which allows hESCs going differentiation (53). We found that a number of AS events of the ATM/ATR pathway are shared by multiple lineages (Supplementary Figure S12b). *BARD1* (BRCA1 associated RING domain 1) is the only gene that shared by all lineages, suggesting its key role in determining cell fate. We observed the AS event of BARD1 involving the exon 3 and 4 for all differentiation lineages (Figure 7a). The stem cells prefer to using exon 3, while differentiated cells prefer to exon 4, with a differential PSI (ΔPSI) of exon 4 at 0.218 in average (Figure 7b). This preference of exon usage is also observed in an additional independent dataset from Roadmap (46,47) and ENCODE (48,49) projects (Supplementary Dataset D2, method M5, and Figure S14a).

**Figure 7.**
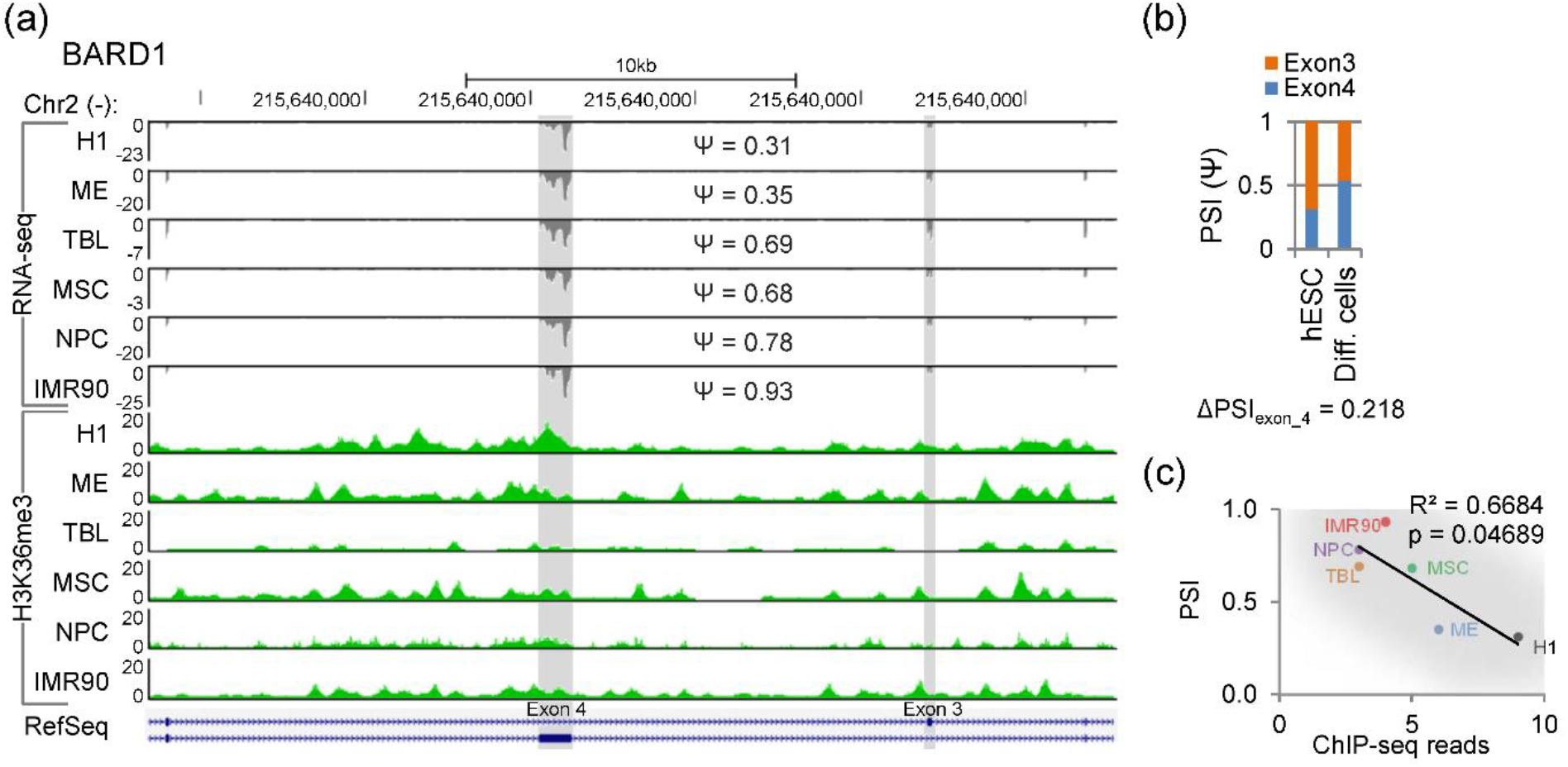
BARD1 is alternatively spliced upon hESCs differentiation and links H3K36me3 to cell fate decision through the ATM/ATR pathway. **(a)** The genome browser view of the AS event of BARD1 and surrounding H3K36me3 signals. Ψ, the PSI of exon 4. **(b)** The isoform switches between hESCs and differentiated cells, shown as the PSIs of exons 3 and 4. **(c)** The inclusion levels of exon 4 are significantly of negative association with the surrounding H3K36me3 signals. p, the p-value, and R^2^, the fitness of linear regression.

In addition, DeepCode highlights the importance of H3K36me3 around the 3’ splice site of exon 4. We observed significantly negative correlation between inclusion levels of exon 4 and the surrounding H3K36me3 signals in hESCs and differentiated cells (Figure 7c). Together with the previous report that chromatin-adaptor MRG15/PTBP1 endows negative regulation of H3K36me3 on AS (26) and the report showing that *BARD1* is a splicing target of PTBP1 (54), the AS of *BARD1* is suggestively regulated by H3K36me3 through the MRG15/PTBP1 adaptor. These results suggest the presence of a combinational mechanism involving HMs and AS, wherein HMs facilitate the PSD and cell fate commitment by impacting on the AS of key components of the ATM/ATR pathway (Supplementary Figures S12c). The underlying mechanisms of these findings are discussed later and hold the promise to drive further experimental studies on the involvements of HMs in cell fate decision via determining the transcripts diversity in addition to their abundance.

## DISCUSSION

Deep learning is now one of the most active fields in artificial intelligence (AI) and has been shown the state-of-the-art performance in image and speech recognition (33-35), natural language understanding (arXiv preprint arXiv:1409.0473 and arXiv:1603.01417), and most recently, in computational biology (12,36-39). Specifically, deep neural networks have been successfully applied to predict splicing activity based only on RNA sequences (12,55). However, genomic features alone limit the regulatory integrity and accuracy in modelling the AS due to the co-transcriptional occurrence and multi-level regulations from chromatin (epigenetics) to RNAs *per se* (genetics). To overcome this limitation, we implemented a multi-label deep neural network (DeepCode) to decode the AS, especially taking epigenetic properties into account (Figure 1). We found that DeepCode significantly improves the performance in predicting splicing patterns during human ESC (hESC) differentiation. In addition, DeepCode highlights the epigenetic mechanisms of AS regulation and reveals that epigenomic features are superior to the genomic ones and pay dominant roles in decoding the AS. Moreover, DeepCode enables extension of this work to other areas beyond the embryonic development, such as cancer research and precision medicine, to widen our knowledge in broader fields (Figure 1c).

Although epigenomic signatures, such as HMs, are mainly enriched in promoters and other intergenic regions, it has become increasingly clear that they are also present in the gene body, especially in exon regions. This indicates a potential link between epigenetic regulation and the predominantly co-transcriptional occurrence of AS. So far, H3K36me3 (26,27), H3K4me3 (24), H3K9me3 (25), and H3acetyl (28) have been revealed to regulate AS, either via the chromatin-adapter systems or by altering Pol II elongation rate. Here, using deep learning approach, we pinpointed the extent to which the HMs are predictive for AS by integrative learning both transcriptome and epigenome data during hESC differentiation. Especially, we revealed high importance of the associations between HMs and AS, including previously reported and unreported ones. H3K36me3 and H3K79me1 were ranked as the top two important HMs in decoding AS events. It is reasonable since H3K36me3 is a mark for actively expressed genomes (56), and has been reported to regulate AS via two chromatin-adaptor systems (26,27); while H3K79me1 has been reported to be highly correlated with splicing (21). Besides, we revealed two more HMs (H3K4me2 and H2BK5ac) that are implied to be related with both MXE and SE events. The H2BK5ac has be linked to active promoters and gene expressions (57,58); H3K4me2 is a mark for both promoters and enhancers (59) and defines the most TF binding regions (60). Both of them have not been reported to associate with or function in AS regulations. Our findings will drive the further studies on these unreported HMs in AS regulations, which are beyond the scope of our current work. Moreover, we assembled the top important HMs and genetic motifs into an ‘extended splicing code’, which gives better performance in decoding splicing and holds the promise for deeper insights into AS regulations.

Additionally, DeepCode also sheds light on understanding of the contribution of HMs to cell fate decision via determining not only what parts of the genome are expressed, but also how they are spliced (23). We found that the alternatively spliced ATM/ATR pathway plays crucial roles in hESC differentiation (Supplementary Figures S12a and S13). Especially, the AS of *BARD1* conservatively occurs in all of the investigated differentiation lineages and is putatively associated with cell fate decision through a process called pluripotency state dissolution (PSD) (Supplementary Figure S12) (53). *BARD1* plays a central role in the control of the cell cycle in response to DNA damage and has also been reported to stabilize the differentiation state (61). BARD1 is specifically homologous to BRCA1 within the conserved RING finger domain at the N-terminus (residues 50-87) (62) and two tandem BRCA1 carboxyl-terminal (BRCT) domains at its C-terminus (residues 560-777) (Supplementary Figure S12c) (63). BRCA1 and BARD1 can form either homodimers or more stable heterodimers via their RING fingers (64), and play key roles in ATM/ATR pathway for DNA repair and cell cycle transition, which are all related to cell fate decision through the PSD (Supplementary Figure S12a) (53). The AS of *BARD1* occurred during hESC differentiation results in two isoforms by mutually including exon 4 and exon 3. The isoform including exon 3 codes the canonic BARD1 protein and is highly expressed in ESCs, which is crucial for the activation of ATM/ATR pathway and then benefits the maintenance of pluripotency state (self-renewal, Supplementary Figure S12c). Whereas, the isoform including exon 4 is a non-coding RNA and undergoes nonsense-mediated decay (NMD) in differentiated cells, which attenuates the activity of ATM/ATR pathway and allows the PSD and further differentiation (Supplementary Figure S12c).

Although the bidirectional communication between H3K36me3 and splicing has been reported (65) and may weaken the regulating hypotheses from H3K36me3 to AS of *BARD1*, the enhancement of H3K36me3 by splicing can only deduce the positive correlations between these two processes. Therefore, our observation of negative correlation putatively suggests a unidirectional regulation from H3K36me3 to AS of *BARD1* (Figure 7c). Based on the previously reports, two chromatin-adaptor complexes, MRG15/PTBP1 (26) and PSIP1/SRSF1 (27), endow negative and positive regulations of H3K36me3 on AS, respectively. Since the observed negative correlation (Figure 7c), it is suggestive that the AS of *BARD1* is epigenetically regulated by H3K36me3 through the MRG15/PTBP1 adaptor system. Based on the independent datasets (Supplementary Dataset D2) from Roadmap/ENCODE projects, the negative correlations between the inclusion level of exon4 and the expressions of MRG15/PTBP1 rather the PSIP1/SRSF1 also implied this regulation hypothesis (Supplementary Figures S14b-e).

Altogether, we implemented a deep learning approach in deciphering a more comprehensive splicing code, DeepCode, integrating both genetic and epigenetic features. Especially, DeepCode revealed a novel candidate mechanism linking histone modifications to ESC fate decision. Besides its outstanding performance in embryonic development, DeepCode is designed for its scalability in data types and extensibility in applications, thus holds the potential applications in a variety of biological and biomedical fields (Figure 1c).

## SUPPLEMENTARY DATA

Supplementary Data are available at NAR online: Supplementary Figures S1-S14, Supplementary Methods M1-M5, Supplementary Datasets D1-D2, and Supplementary References [1-16].

## ACKNOWLEDGEMENT

The authors thank the editors and reviewers for their valuable comments and suggestions.

